# Modeling protected species distributions and habitats to inform siting and management of pioneering ocean industries: A case study for Gulf of Mexico aquaculture

**DOI:** 10.1101/2022.04.07.487536

**Authors:** Nicholas A. Farmer, Jessica R. Powell, James A. Morris, Melissa S. Soldevilla, Lisa C. Wickliffe, Jonathan A. Jossart, Jonathan K. MacKay, Alyssa L. Randall, Gretchen E. Bath, Penny Ruvelas, Laura Gray, Jennifer Lee, Wendy Piniak, Lance Garrison, Robert Hardy, Kristen M. Hart, Chris Sasso, Lesley Stokes, Kenneth L. Riley

## Abstract

Marine Spatial Planning (MSP) provides a process that uses spatial data and models to evaluate environmental, social, economic, cultural, and management trade-offs when siting ocean industries. Aquaculture is the fastest-growing food sector in the world. The U.S. has substantial opportunity for offshore aquaculture development given the size of its exclusive economic zone, habitat diversity, and variety of candidate species for cultivation. However, many protected species rely upon habitats that overlap with promising aquaculture areas. Siting surveys, farm construction, operations, and decommissioning can alter the habitat and behavior of animals in the vicinity of these activities. Vessel activity, underwater noise, and physical interactions between protected species and farms can potentially increase the risk of injury or cause direct mortality. In 2020, the U.S. Gulf of Mexico was identified as one of the first regions to be evaluated for offshore aquaculture opportunities as directed by a Presidential Executive Order. We developed a generalized scoring model for protected species data layers that captures vulnerability using species conservation status and demographic information. We applied this approach to data layers for eight species listed under the Endangered Species Act, including five species of sea turtles, Rice’s Whale, Smalltooth Sawfish, and Giant Manta Ray. We evaluated several methods for scoring (e.g., arithmetic mean, geometric mean, product, lowest scoring layer) and created a combined protected species data layer that was used within a multi-criteria decision-making modeling framework for MSP. The product approach for scoring provided the most logical ordering of and the greatest contrast in site suitability scores. This approach provides a transparent and repeatable method to identify aquaculture site alternatives with the least conflict with protected species. These modeling methods are transferable to other regions, to other sensitive or protected species, and for spatial planning for other ocean-uses.

## Introduction

Marine Spatial Planning (MSP) provides a solutions-focused pathway for ocean planning through the use of spatial data and models to evaluate interactions across multiple spatial and temporal scales. An MSP approach provides opportunities to evaluate tradeoffs among environmental, social, economic, cultural, and management considerations to inform how and where ocean and coastal industries can be sited. Ecosystem-level MSP (i.e., planning at the regional or ocean basin scale) presents unique challenges where expanded datasets and broader scale determinations are required and data are often limited.

MSP approaches are being implemented throughout the world to determine space appropriate for pioneering ocean industries (e.g., wind energy, aquaculture) (Lester et al. 2018; Spijkerboer et al. 2020). Finding the most suitable ocean space for emerging industries is challenging, given historic and current ocean uses and the potential for disrupting established industries that could have global socioeconomic impacts. Introducing new or additional anthropogenic stressors to an ecosystem requires appropriate consideration of cumulative impacts from industries, especially in the context of environmental change (e.g., climate change). Given the demand for ocean-based food and energy production, it is imperative that approaches developed to find suitable space rely on MSP in order for regulators and scientists to make informed decisions about potential ecosystem or resource impacts.

Aquaculture is the fastest growing food sector in the world; however, only seven percent of the U.S. seafood supply comes from aquaculture (FAO 2020). The U.S. ranks among the top nations in the world in opportunity for offshore aquaculture development given the size of the U.S. economic exclusive zone (EEZ; 3.4 million square miles), the diversity of habitats (polar to tropical), and variety of candidate cultivation species. Innovative farm design and engineering provide aquaculture gear and cultivation approaches that can withstand dynamic offshore conditions and increase production capacity (Langan 2012; Holm et al. 2017; Chu et al. 2020).

Aquaculture development in sensitive marine habitats carries inherent risks depending upon aquaculture type and operational standards with naturally-occurring marine species. Vulnerable anadromous and marine protected species, including marine mammals, sea turtles, and fishes, are reliant upon marine habitats (e.g., surface, mid-water, and benthic environments) for survival and success. Aquaculture development impacts specific to protected resources can vary across a range of activities including environmental and cultural resource surveys, construction, operation and management, and farm decommissioning. Potential impacts can include attraction to farms or displacement from critical habitats which can result in changes to distribution, behaviors, or social structures (Clement 2013; Price et al. 2017; Heinrich et al. 2019). Physical interactions with gear, vessel traffic, noise, and light pollution can also result in injuries or mortalities (Hamelin et al. 2017; Price et al. 2017; Callier et al. 2018). When farms are sited and managed properly, impacts to protected resources can be minimized by avoiding sensitive areas or time periods and using best management practices for construction and operation. As the aquaculture industry continues to grow worldwide, cumulative impacts to protected species are an important siting consideration because farms could be developed in areas with existing anthropogenic threats (NASEM 2017).

U.S. Presidential Executive Order 13921 (E.O.) (May 7, 2020) called for the expansion of sustainable seafood production in the U.S. The directive required the Secretary of Commerce, in consultation with relevant federal agencies, to identify Aquaculture Opportunity Areas (AOA) suitable for potential commercial offshore aquaculture development. Under the E.O., AOAs should use comprehensive data acquisition, novel modeling methodologies, and public stakeholder engagement to support environmental, economic, and social sustainability; and minimize unnecessary resource use conflicts. To support the E.O., NOAA’s National Centers for Coastal Ocean Science (NCCOS) commenced an MSP study to identify suitable AOA options in federal waters of the U.S. Gulf of Mexico and southern California (Riley et al. 2021; Morris et al. 2021).

Under this E.O. directive and following the legal mandates of the U.S. Endangered Species Act (ESA) (16 USC. 1531 et seq.) and Marine Mammal Protection Act (MMPA) (16 USC. § 1361) to avoid or minimize effects to protected species, we developed a novel approach to integrate pertinent marine protected species data into the AOA MSP model. In each region, AOA options that minimize potential interactions with vulnerable species and critical habitats for reproduction, feeding, migration, and residency were identified. Although specific to planning for AOAs, this standardized integration of transient protected species data into a multi-criteria decision-making MSP modeling framework is transferable to assess the potential risk of other protected species and ocean-use conflicts. The main goal of this study is to provide generalized guidance for avoiding development conflicts with protected species in the marine environment. Here, a case study is provided for an MSP modeling process for AOAs in the U.S. Gulf of Mexico.

## Methods

### Developing generic scoring for protected resources data layer

We developed a generalized scoring system to measure protected species vulnerability based on species status under the ESA or MMPA, population size, and population trajectory to inform relative risk in spatial modeling (**Table 1**). Because specific aquaculture activities are not identified upfront with the creation of an AOA, a generalized approach was desired that could universally provide a general, comparable measure of protected species risk across aquaculture types and regions. Under this generalized system, scores for MMPA and ESA-listed species data layers range from 0.1 (most vulnerable species, based on their biological status) to 0.8 (least vulnerable species) based on the best available data for each protected species considered (**Table 1**). Species and MMPA stocks are ranked according to factors that are more or less likely to affect their ability to withstand mortality, serious injury, or other impacts to the species’ ability to survive and recover. A stock is defined by the MMPA as a group of marine mammals of the same species or smaller taxa in a common spatial arrangement that interbreed when mature. A generalized score of 0.5 was also reserved for layers or areas within layers to reflect types of data needing further consideration during the site characterization process. The generalized score is the standard score used in the spatial model for data where suitability or compatibility of a location for potential aquaculture activities is uncertain. A score of 1 reflects an area with no protected species conflict whereas a score of 0 reflects an area that is unsuitable for an aquaculture farm given protected species vulnerability. A generalized scoring table reflective of relative differences in statutory protection, status, and trend is consistent with U.S. Congressional reporting through the Government Performance and Results Act (GPRA) process (5 U.S.C. 306 Chapter 3).

**Table 1.** A generalized scoring system for endangered and threatened species data layers.

| Status | Trend | Converted scores for model |
| --- | --- | --- |
| **Endangered** | declining, small population* or both | 0.10 |
| **Endangered** | stable or unknown | 0.20 |
| **Endangered** | increasing | 0.30 |
| **Threatened** | declining or unknown | 0.40 |
| **Threatened** | stable or increasing | 0.50 |
| **MMPA Strategic** | declining or unknown | 0.60 |
| **MMPA listed** | small population* | 0.70 |
| **MMPA listed** | large population | 0.80 |

*\*Small population equates to populations of 500 individuals or less (Franklin 1980).*

### Combining Protected Species Data Layers

To determine the most conservative approach providing the highest contrast in the AOA model, four mathematical approaches to combine protected species data layers were evaluated: 1)

Geometric Mean (*g*) (Equation 1), 2) Arithmetic Mean (*µ*) (Equation 2), 3) Product (*ρ*) (Equation 3), and 4) Lowest Scoring Layer (*l*) (Equation 4). These are computed as follows:

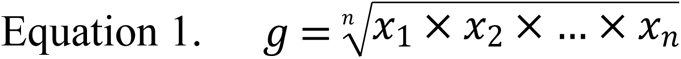

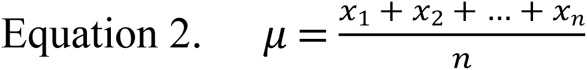

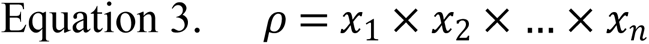

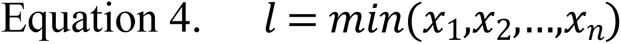

where *x_n_* represents cell-specific scores based on species status and trend (**Table 1**). These methods were applied to hypothetical protected species data layers to evaluate their performance with regards to directionality and spread of combined scores. We considered a geographically- constricted, endangered species with a declining, small population (score = 0.1); a broadly-distributed declining threatened species (score = 0.4); and a broadly-distributed marine mammal with a large population (score = 0.8).

### Case Study: Gulf of Mexico Aquaculture

NOAA Fisheries has been tasked with identifying AOAs over seven or more years using the best-available science to identify areas suitable for domestic aquaculture production, while minimizing impacts to NOAA-trust resources. The first phase of the AOA process is the development of a public-facing Atlas that visually represents the relative suitability of different locations for potential aquaculture activities. The study areas for the Gulf of Mexico potential AOA are in federal waters between 50–150 m depth and delineated using an ecosystem-based approach that results in four separate study areas (West, Central, East, and Southeast Gulf of Mexico study areas). The four subregions were evaluated separately to promote geographic representation. The Atlas study area was derived from the U.S. Coastal Relief Model bathymetry raster (NOAA NCEI 2020). A polygon was created encompassing depths from 50–150 m, bounded in the west by the U.S. EEZ, and in the east at -80.17° longitude.

We evaluated the best scientific information available for the distributions of vulnerable ESA-listed species within the proposed Gulf of Mexico Atlas study areas (Hayes et al. 2021). Individual spatial data layers for the species listed in Table 2 were developed, assigned scores (**Table 2**), and then combined to create a comprehensive protected species layer for the Atlas study area. ESA-listed Sperm Whales were not included because they typically occur in waters deeper than the Atlas study areas (Hayes et al. 2021). Gulf Sturgeon was not evaluated, as the species occurs primarily inshore of the 50-m depth contour. Similarly, Oceanic Whitetip Shark was not considered, as the species occurs primarily offshore of the 150-m depth contour, although several shark species have been observed at aquaculture sites (Papastamatiou et al. 2010; Blue Ocean Mariculture LLC 2014), with records of Oceanic Whitetip Sharks at offshore (2000 m depth) Hawaiian aquaculture cages (Y. Papastamatiou, M. Hutchinson, pers. comm.). Marine mammals protected solely under the MMPA were not included in the data layer because insufficient time and resources were available to obtain and evaluate their distributions relative to the proposed Atlas study areas.

**Table 2.** The eight ESA-listed species known to occur within the Gulf of Mexico Atlas study areas and their suitability scores, as determined by species status and trend. Note: ESA-listed corals are not included because areas containing corals are scored as ‘0’ (not suitable for aquaculture activities) for the Atlas.

| *Common name* | *Species name* | *Status and Trend* | *Score* |
| --- | --- | --- | --- |
| *Rice’s Whale*<sup>1</sup> | *Balaenoptera ricei* | Endangered, small and declining | 0.1 |
| *Leatherback Sea Turtle* | *Dermochelys coriacea* | Endangered, declining | 0.1 |
| *Kemp’s Ridley Sea Turtle* | *Lepidochelys kempii* | Endangered, unknown | 0.2 |
| *Hawksbill Sea Turtle* | *Eretmochelys imbricata* | Endangered, unknown | 0.2 |
| *Smalltooth Sawfish, U.S. DPS* | *Pristis pectinata* | Endangered, increasing | 0.3 |
| *Giant Manta Ray* | *Manta birostris* | Threatened, declining | 0.4 |
| *Loggerhead Sea Turtle, Northwest Atlantic Ocean DPS* | *Caretta caretta* | Threatened, unknown | 0.4 |
| *Green Sea Turtle, North Atlantic DPS* | *Chelonia mydas* | Threatened, increasing | 0.5 |

<sup>1</sup>Formerly Gulf of Mexico Bryde’s Whale (*Balaenoptera edeni*)

Submodels in the AOA relative suitability model provide a consistent, categorical structure for organizing complex and dynamic ocean data within the planning exercise (Lightsom et al. 2015). Equally-weighted submodels included 1) National Security; 2) Industry, Navigation, and Transportation; 3) Commercial Fishing and Aquaculture; 4) Natural and Cultural Resources; and 5) Constraints (Riley et al. 2021). The Natural and Cultural Resources submodel included the protected species data layer. The Constraints submodel included activities and occupied areas unsuitable for aquaculture siting (e.g., military zones, coral, and hardbottom) and were eliminated before calculating the geometric mean and creating a cumulative suitability score for each study area (i.e., West, Central, East, Southeast). A Localized Index of Spatial Association (LISA) was then implemented on the cumulative suitability output for all study areas to determine statistically significant clusters (p < 0.05), or the grid cells with the highest suitability relative to others, within a given study area. Within the most suitable clusters in each study area, a two-stage precision siting model was then used to identify and rank multiple AOA options within each cluster and then, secondarily, among clusters of the study areas, which were then characterized and used to provide specific details of preliminary options for AOAs (**Figure 1**).

**Figure 1.**
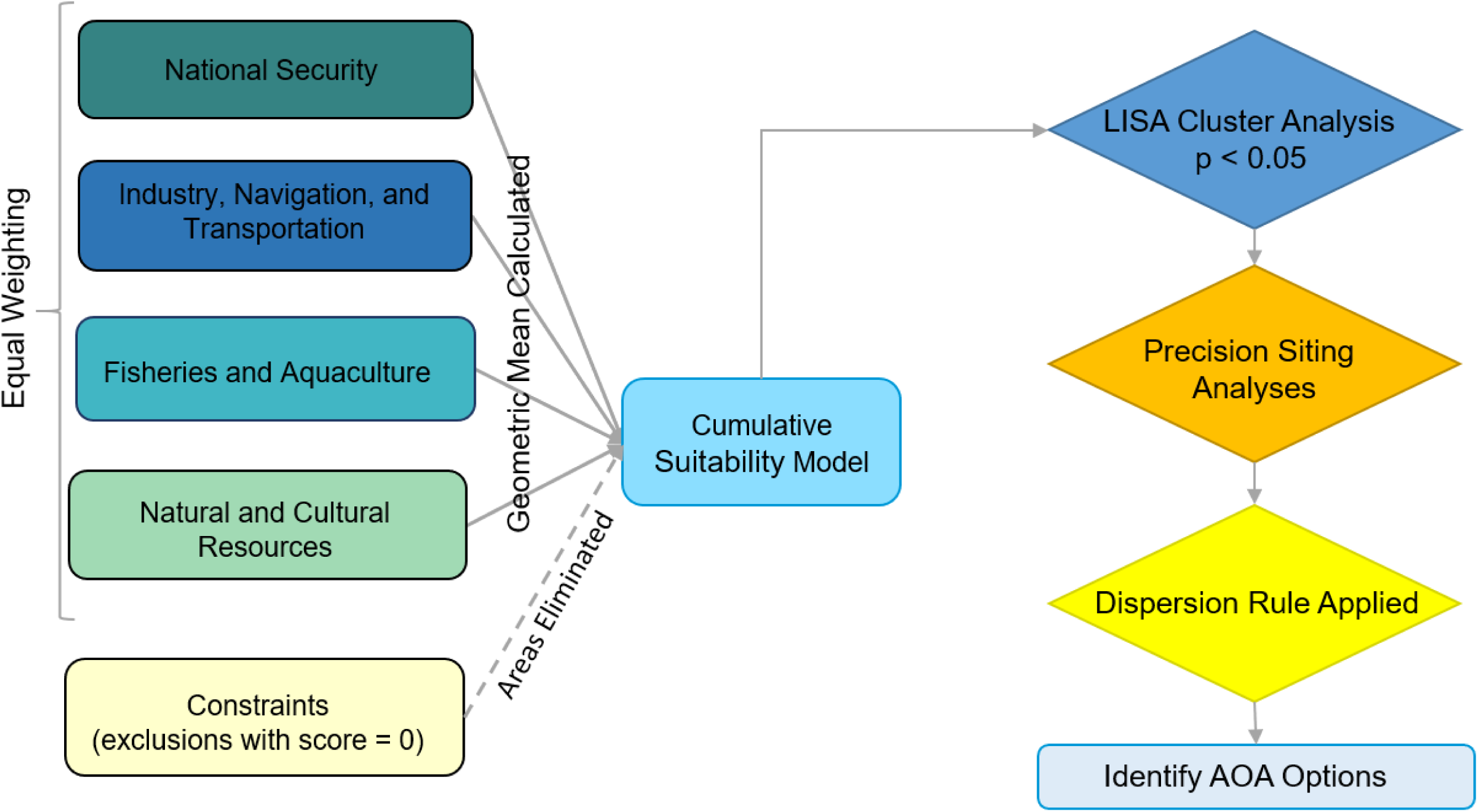
Flow chart illustrating the Aquaculture Opportunity Area (AOA) Atlas model process used to identify final AOA options. Note that equal relative weighting is imposed when combining the submodels, including the Natural Resources submodel, which contains the protected species data layer.

An equally-weighted site suitability model was utilized for the AOA study because multiple types of aquaculture are proposed making it difficult to compare between potential spatial conflicts. For example, bottom culture techniques and gear may have different requirements than surface culture techniques. Because the final suitability scores compare different types of activities, the overall score does not have an empirical meaning; however, the relative rank of the scores is important to determining which options are more suitable. Therefore, it is important that the ranked scores of the areas in the protected species layer of the Natural Resources submodel are reflective of where managers are more or less concerned with regards to the vulnerability of protected species.

### Rice’s Whale

The Rice’s Whale is the only species of large whale indigenous to the United States (Rosel et al. 2021). The population is estimated at fewer than 100 individuals, with mean estimates of <50 individuals remaining (Rosel et al. 2021). The species was listed under the ESA in 2019 and is exposed to a number of threats in the highly industrialized northern Gulf of Mexico, including vessel strikes, interactions with commercial fisheries, and exposure to industrial noise (Soldevilla et al. 2021; Rosel et al. 2014). As a long-lived marine mammal with low reproduction rates and a very small population size, the loss of a single individual could drive the species to extinction (Franklin 1980; Rosenfeld 2014). Available data on Rice’s whale distribution in the northern Gulf of Mexico support a core habitat (**Figure 2A**, yellow) and an extended habitat (Figure 2A, green). The core habitat (**Figure 2A**, yellow) is based on the best available tag and sightings data available as of June 6, 2019 (Rosel et al. 2020; Rosel and Garrison 2022). A convex hull polygon was developed around all recorded northeastern Gulf of Mexico baleen whale (Rice’s Whale, Rice’s/Sei, Rice’s/Sei/Fin) sighting locations (n=119) from surveys from 1998–2018, telemetry tag locations (n=52) from a single Rice’s Whale tagged in 2010, and Acousonde tag locations (n=41) for a whale tagged in 2015 (Soldevilla et al. 2017; Rosel et al. 2020; Rosel and Garrison 2022). The polygon was buffered by 30 km to account for the 10 km strip width of surveys and the approximately 20 km median daily range of movements from satellite tagged animals (Rosel and Garrison 2022). This buffered polygon was trimmed on the western-side to the 410 m isobath, determined based on the current deepest known sighting of 408 m (Soldevilla et al. 2017; Rosel and Garrison 2022).

**Figure 2.**
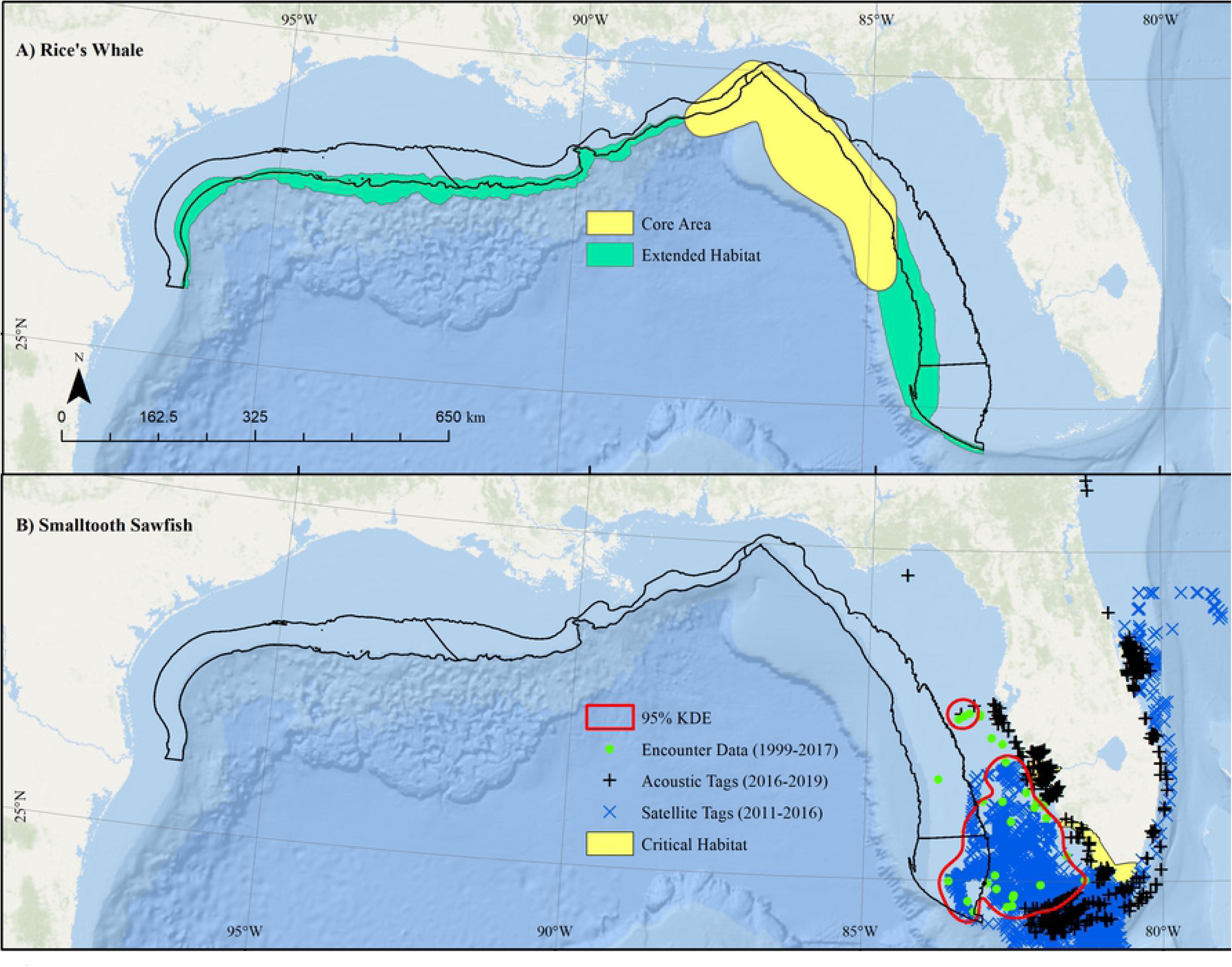
Distribution of A) Rice’s Whale and B) Smalltooth Sawfish relative to proposed aquaculture opportunity atlas study areas (black). Rice’s Whale distribution polygons based on sightings (yellow “core area”) and sightings coupled with passive acoustic monitoring (green “extended habitat”). Sawfish high use areas (red line) based on 95% kernel density estimate for observations fit to pooled data from Sawfish Encounter Database (1999–2017; 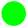), acoustic tag detections (2016-2019; **+**), and positioning estimates from satellite tags (2011–2016; 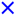). Note Sawfish critical habitat (yellow) did not overlap the proposed aquaculture area. Map developed in ESRI ArcGIS 10.8.1. Basemap from Esri Ocean Basemap and its partners. Projection: UTM NAD83 Zone 17N.

The extended habitat (green) in **Figure 2A** represents the distribution of Rice’s Whale beyond the core habitat in the western and eastern Gulf of Mexico between the 100–400 m isobaths, inferred from sightings data, long-term passive acoustic monitoring (PAM), and habitat suitability modeling (Garrison 2021; M. Soldevilla, unpublished data). The supporting data for the extended habitat include 1) all recorded Gulf of Mexico baleen whale sighting locations from surveys from 1998–2018 and 2) detections of baleen whale calls in one year of acoustic recordings from passive acoustic moorings deployed in areas of predicted Rice’s Whale habitat along the shelf-break from near DeSoto Canyon in the east to offshore of the Flower Garden Banks National Marine Sanctuary (FGBNMS) in the west during 2016–2017 (M. Soldevilla, unpublished data). In August 2017, a genetically verified sighting of a Rice’s Whale occurred along the shelf break offshore of Corpus Cristi, Texas. Additionally, stereotypical Rice’s Whale long-moan type calls (Rice et al. 2014, Soldevilla et al., in review) were detected at three of four northcentral/northwestern Gulf sites (Soldevilla et al., in review). Rice’s Whale calls were persistently detected throughout the year, with no evidence of seasonality, on 16% of days at the westernmost site near the FGBNMS as compared to 1% of days at the central site off Eugene Isle, supporting the year-round importance of the extended habitat area (Soldevilla et al., in review). By comparison, long-moan calls are detected more frequently, on 90–100% of days, in the eastern Gulf of Mexico core habitat (Rice et al. 2014), though effects of site-specific sound propagation, ambient noise levels, and whale calling rates make it difficult to directly infer site-specific density of animals present from acoustic presence data. This highlights the importance of two distinct habitat regions until the distribution and density of this species is better understood. In contrast, a passive acoustic mooring deployed at 800 m depth offshore of Alabama for two years had no detections of Rice’s Whale calls (M. Soldevilla, unpublished data), suggesting limited occurrence in deeper waters.

This understanding of Rice’s Whale distribution is further supported by habitat preference modeling results. Preliminary spatial density modeling efforts for Rice’s Whale based on sightings data identified a relatively high density area ranging from shelf-edge Alabama to southwest Florida, with further suitable habitat in a more narrow shelf-edge strip extending to central Texas to the west and the Florida Keys to the east (Roberts et al. 2016). This model was based on general habitat features and may not have captured the biological and physical conditions required to support the Rice’s Whale population.

On 14 surveys from 2003–2019, 154 Rice’s Whale groups were sighted using a combination of directed surveys and multispecies line transect surveys. A generalized additive model (GAM) was used to evaluate the environmental drivers of Bryde’s Whale distribution. Variables considered included depth, SST, surface and bottom salinity, SSH, velocity, log chlorophyll-a (Chl-a), and bottom temperature. The selected model included depth, bottom temperature, log Chl-a, and SSH. This model was used to predict Rice’s Whale distribution both within the core habitat and elsewhere in the broader Gulf of Mexico. The model showed good fits to sightings data and low uncertainty within the sampled region. It showed higher uncertainty in deep and shallow waters, but had very low predicted occurrence in those habitats. Similar to PAM findings above, the modeling process, which did not consider the PAM data, identified a probable distribution along the Gulf of Mexico shelf break concentrated in the core area but extending in a narrow band contained within the 100–400 m depth contours following preferred bottom temperatures along the shelf break throughout the Gulf of Mexico. Therefore, the 100–400 m depth contour was considered an appropriate proxy for the model results. The increased concentration of Rice’s Whale in the core habitat appeared to be explained by notably higher summer Chl-a concentrations in that area as compared to other regions with suitable bottom temperatures. This area is characterized by seasonal advection of low salinity, high productivity surface waters, leading to persistent upwelling driven by both local processes (winds) and intrusion of Loop Current features. Rice’s Whales are most commonly observed in the mixing area, characterized by intermediate (non-oceanic) Chl-*a* concentrations, intermediate bottom temperatures, and high salinity bottom water at the boundary between coastal and deep oceanic waters. Other regions in the Gulf have similar bottom temperatures at the shelf-break, but less surface productivity, which may partially explain the less frequent observations of the species in those areas. The areas west of the core distribution are also characterized by much higher levels of shipping activity and noise associated with oil and gas exploration, both of which have been identified as threats to the species and implicated in the possible contraction of their geographic range (Rosel et al. 2016).

Sightings data, habitat modeling, and PAM data all suggest a core area in the northeastern Gulf and an extended habitat area across the northern Gulf between 100–400 m depth contour with less frequent but year-round occurrences. We assigned a score of 0.1 to the union of the two Rice’s Whale layers.

### Smalltooth Sawfish, U.S. DPS

Smalltooth Sawfish are rays with a long, flat rostrum edged with teeth. Smalltooth Sawfish populations declined dramatically during the second half of the 20th century due to habitat loss associated with coastal development and accidental capture in fisheries. They were listed as endangered under the ESA in 2003. Smalltooth Sawfish location data were obtained from three point sources: 1) US Sawfish Recovery Encounter Database (Simpfendorfer and Wiley 2006; International Sawfish Encounter Database 2021), 2) Acoustic tag data (Graham et al. 2021; Graham et al. 2022), and 3) Satellite tag data (Carlson et al. 2014) (**Figure 2B**). The US Sawfish Recovery Encounter Database provides data on sawfish observations from 1999–2017; additional data are continually added. The U.S. Smalltooth Sawfish Recovery Team (Smalltooth Sawfish Recovery Implementation Team 2021) manages the most updated version of domestic sawfish encounter records and shares these with other databases including the International Sawfish Encounter Database (ISED), curated by the Florida Program for Shark Research (FPSR) at the Florida Museum of Natural History on the University of Florida campus (International Sawfish Encounter Database 2021). Information from verified sawfish encounter reports is entered into the database and used for monitoring sawfish populations. This information assists in the evaluation of species abundance and range, helping to estimate the population sizes and also to identify habitat preferences. This type of information is vital for monitoring the recovery of worldwide sawfish populations and greatly assists in conservation efforts.

Between May 2016 and April 2019, researchers have used passive acoustic telemetry and 3 large data sharing networks of receivers to track movements of 43 large juvenile and adult Smalltooth Sawfish. During this study, 24 females and 19 males were implanted with transmitters with estimated 4- or 10-year battery lives. These tagged individuals were detected off the southeastern United States via 461 receivers ranging from off the coast of Brunswick, Georgia, to the lower Florida Keys, and along the Gulf coast to Apalachee Bay, Florida. Seasonal migrations were undertaken by 58% (43% mature; 57% immature) of the tagged individuals, with the remainder being apparent residents of their tagging locations (**Figure 2B**). Tagged juvenile and adult sawfish of both sexes migrated, which indicates that neither sex nor maturity is a predictor of whether a sawfish will migrate or not. Although both coasts of Florida were used for migration, most individuals consistently used the same coast when they migrated. The areas surrounding Boca Grande, Cape Canaveral, and the lower Florida Keys were the most heavily visited sites.

Since 2011, members of the Smalltooth Sawfish Recovery Team have been satellite-tagging juvenile and adult sawfish to track broad-scale movements (Carlson et al. 2014; Graham et al. 2021; Graham et al. 2022). Maximum likelihood positioning estimates generated by Wildlife Computers GPE3 positioning software for 15 satellite-tagged sawfish were provided by Jasmin Graham (Florida State University, 2021).

Smalltooth Sawfish point data derived from the three aforementioned sources were merged into a single GIS dataset and filtered to include only locations in the Gulf of Mexico EEZ. A 95% Kernel Density Estimate (KDE) was generated to encompass a Smalltooth Sawfish high-use area using the *kernelUD* function in the ‘adeHabitat’ package in *R v4.1.2* (**Figure 2B**, yellow polygons). All cells intersecting this 95% KDE spatial extent received a score of 0.3, while all other cells received a score of 1 (i.e., high suitability) for Smalltooth Sawfish.

### Sea Turtles

All species of sea turtles in U.S. waters are listed under the ESA, primarily due to global threats including incidental capture in fishing gear (bycatch), illegal harvest, habitat loss, and egg collection for human consumption. Leatherback, Green, Hawksbill, Kemp’s Ridley, and Loggerhead Sea Turtle habitat use and distribution overlap with the Gulf of Mexico and Atlas study areas. Leatherback Sea Turtles are the largest turtles in the world, with the widest global distribution of any reptile. They are the only species of sea turtle that lacks a hard shell and forage throughout the water column on jellyfish and salps. Green Sea Turtles inhabit sub- tropical and temperate regions and are unique among sea turtles in that they are herbivorous, foraging largely on seagrass and algae. Hawksbill Sea Turtles inhabit tropical and sub-tropical waters and forage on sponges following their pelagic stage. Kemp’s Ridley Sea Turtles are found only in the North Atlantic Ocean, primarily in the Gulf of Mexico, and forage mostly on crabs. Loggerhead Sea Turtles inhabit sub-tropical and temperate regions and forage on benthic invertebrates such as mollusks and crabs. We identified available data that could be used to characterize important areas for sea turtles within the study area.

Six sea turtle datasets were obtained: (1) Aerial survey data from the Gulf of Mexico Marine Assessment Program for Protected Species (GoMMAPPS) (three surveys from 2017–2018) and National Resources Damage Assessment (NRDA) (four surveys from 2011–2012) efforts which spanned the northern Gulf of Mexico and covered the aquaculture study area, including observations of 2,253 Loggerheads, 1,209 Kemp’s Ridleys, 276 Greens, and 252 Leatherbacks; (2) Residence area locations from multiple satellite telemetry studies (Hart et al. 2021), including records from 188 Loggerheads, 72 Greens, 42 Hawksbills, and 33 Kemp’s Ridleys; (3) Leatherback distribution data in the northern Gulf of Mexico (Aleksa et al. 2018); (4) Residence area locations from a satellite telemetry study which included 15 adult female Green Sea Turtles (Schroeder B, unpublished data); (5) Sea turtle observations made by fishery observers in the Gulf of Mexico during 2005–early 2020 in the pelagic longline, shrimp, reef fish, gillnet and shark bottom longline fisheries, including observations of 365 incidentally captured Leatherbacks, 180 Loggerheads, 136 Kemp’s Ridleys, 35 Greens, and 1 Hawksbill (SEFSC, unpublished data); and (6) Residence area locations from 81 adult female loggerheads tracked as part of several satellite telemetry studies (Hardy et al. 2014).

These datasets were analyzed in a geographic information system (GIS) and used to produce a final representation of high-use areas (HUAs) within the Atlas study area. First, residence area locations derived from satellite telemetry data were converted to polygons (buffered 18.983 km radius) to produce representations of high-use residence areas consistent with those identified in the literature; specifically, the chosen buffer distance produced polygons that were 1,132 km^2^ which was equivalent to the mean size of loggerhead 90 or 95% utilization distributions reported in previous studies (Foley et al. 2014; Hart et al. 2014; Phillips et al. 2021). Next, separate kernel density (KD) surfaces were created from the aerial survey and fishery observations. For the fishery observations, we used a search radius of 19 km within the KD as was used for the telemetry dataset. We used a KD search radius of 40 km for the observations made during aerial surveys; the increased search radius was used for these data to account for survey transect spacing and sea turtle movements. The resultant density surfaces were classified into quartiles and the upper two quartiles were extracted as HUAs, producing two outputs, one based on aerial survey observation density and another based on fishery observer observation density. Those three polygon outputs were combined with the Atlas polygon to identify portions of the Atlas study area that were HUAs by species for Hawksbills, Loggerheads, Greens, and Kemp’s Ridleys. For Leatherbacks, only the aerial survey and fishery observer data were used to produce HUAs (**Figure 3**). The resultant output encompassed the identified HUAs in Aleksa et al. (2018); thus, the inclusion of telemetry data was not necessary for this species. A recently published Leatherback telemetry study confirmed the HUAs identified using the previous telemetry and observer data (Sasso et al. 2021). Insufficient data were available on Hawksbill turtles to identify HUAs using these methods.

**Figure 3.**
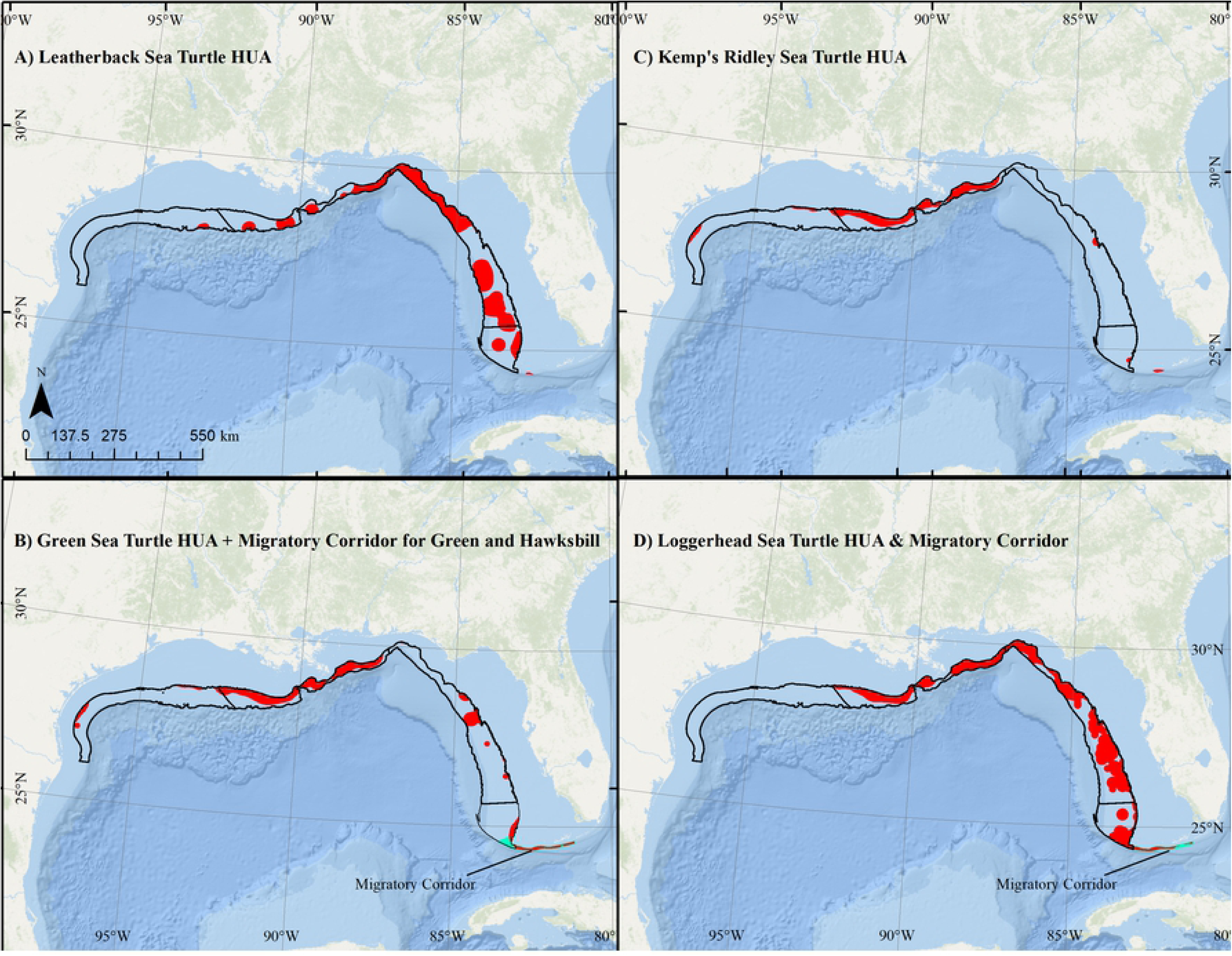
High use areas (HUA; red) and migratory corridors (blue) for A) Leatherback Sea Turtle B) Green and Hawksbill Sea Turtles; C) Kemp’s Ridley Sea Turtle; and D) Loggerhead Sea Turtles relative to proposed aquaculture study areas (black). Note, due to insufficient data, no high use areas were identified for Hawksbill Sea Turtles within the aquaculture study areas. Map developed in ESRI ArcGIS 10.8.1. Basemap from Esri Ocean Basemap and its partners. Projection: UTM NAD83 Zone 17N.

The southern portion of the Atlas study area, near the Florida Keys, was identified as a high-use migratory corridor for Loggerheads, Greens, and Hawksbills based on a review of previous satellite telemetry studies (Foley et al. 2013; Shaver et al. 2016; Iverson et al. 2020; Hart et al. 2021).

### Giant Manta Ray

The Giant Manta Ray (*Manta/Mobula birostris, Manta/Mobula cf. birostris)* is the world’s largest ray species with a maximum wingspan of 8.8 m. They are filter feeders and eat large quantities of zooplankton. The main threat to the Giant Manta Ray is commercial fishing, with the species both targeted and caught as bycatch in a number of global fisheries throughout its range. Few dedicated surveys for Giant Manta Ray exist; however, due to their large size and distinct appearance, they are often observed and recorded during visual aerial surveys that target marine mammals and sea turtles. Farmer et al. (2022) assembled a GIS database of manta sightings from peer-reviewed literature, survey databases, gray literature reports, and anecdotal sources (e.g., social media, press reports, personal communications).

Informal interviews with observers were used to assess the reliability of identification records from surveys and gray literature. Photographs and videos were used to verify the accuracy of species identification from anecdotal sources. Over 5000 Giant Manta Ray sightings were reported in the northwestern Atlantic Ocean and Gulf of Mexico from 1925–2019. Distance-weighting methods were used to account for individual survey effort and species distribution models (SDMs) were fit to a combined dataset generated from 1) Southeast Fisheries Science Center (SEFSC), 2) North Atlantic Right Whale Consortium (NARWC), and 3) New York State Energy Research and Development (NYSERDA) aerial surveys. Generalized additive models (GAMs) were fit to all possible permutations of bathymetry and environmental parameters using the ‘*mgcv’* package in R (Wood and Wood 2015). GAMs were fit with a binomial distribution using a logit link function, with tensor splines limited to 3 knots, such that the resultant SDMs describe the probability of species presence, also termed “habitat suitability” (Brodie et al. 2018) or “habitat preference” (Hazen et al. 2017). Farmer et al. (2022) selected the best-fitting model by lowest AIC and compared it to competing GAM configurations tiered off the best-fitting GAM by excluding non-significant terms in the model summary. The final model was selected by comparing residual deviance explained and predictive power as evaluated through 10-fold internal cross validation and external validation using independent sources (Farmer et al. 2022).

The best GAM fit to combined SEFSC, NARWC, and NYSERDA surveys explained 19% of residual deviance and predicted higher probabilities of observation with warm sea surface temperatures (SST), moderate Z-transformed SST frontal gradients (Front-Z), nearshore and shelf-edge depths, moderate bathymetric slopes, and increasing Chl-a concentrations (Farmer et al. 2022). Measures of predictive utility from internal and external validation were comparable and indicated “excellent” model fits (Hosmer et al. 1988). The Farmer et al. (2022) combined survey model predicted highest probabilities of detection at offshore sloped habitats (e.g., seamounts) and in the nearshore environment off Louisiana at the Mississippi River delta between April to June and again in October. Probability of detection increased at moderate frontal gradients with SSTs between 20–30 °C in both nearshore and shelf-edge environments with moderate slopes and high concentrations of Chl-a. External validation using median Z-score standardized probabilities of observation (Farmer et al. 2017; Heyman et al. 2019) confirmed that SDM model predictions were highly consistent with independent observations of Giant Manta Ray (*t*(4025) =128.01, *p* < 0.0001; Farmer et al. 2022).

To evaluate Giant Manta Ray distribution relative to the Atlas, the final Farmer et al. (2022) combined survey SDM was fit to monthly data from January 2003 to December 2019. The maximum predicted species presence across these 204 months was retained in a final predictive grid for each cell (**Figure 4A**). To provide meaningful contrast to inform the Atlas site selection process, several potential cutoffs were evaluated based on quantiles for maximum probability of presence that would receive the **Table 1** score of 0.4. Because predictions from the Giant Manta Ray SDM are not normally distributed, the median is the most appropriate measure of central tendency. Areas above the median maximum predicted value were designated as HUAs and assigned a score of 0.4 (**Figure 4B**); all other areas received a score of 1 for Giant Manta Ray.

**Figure 4.**
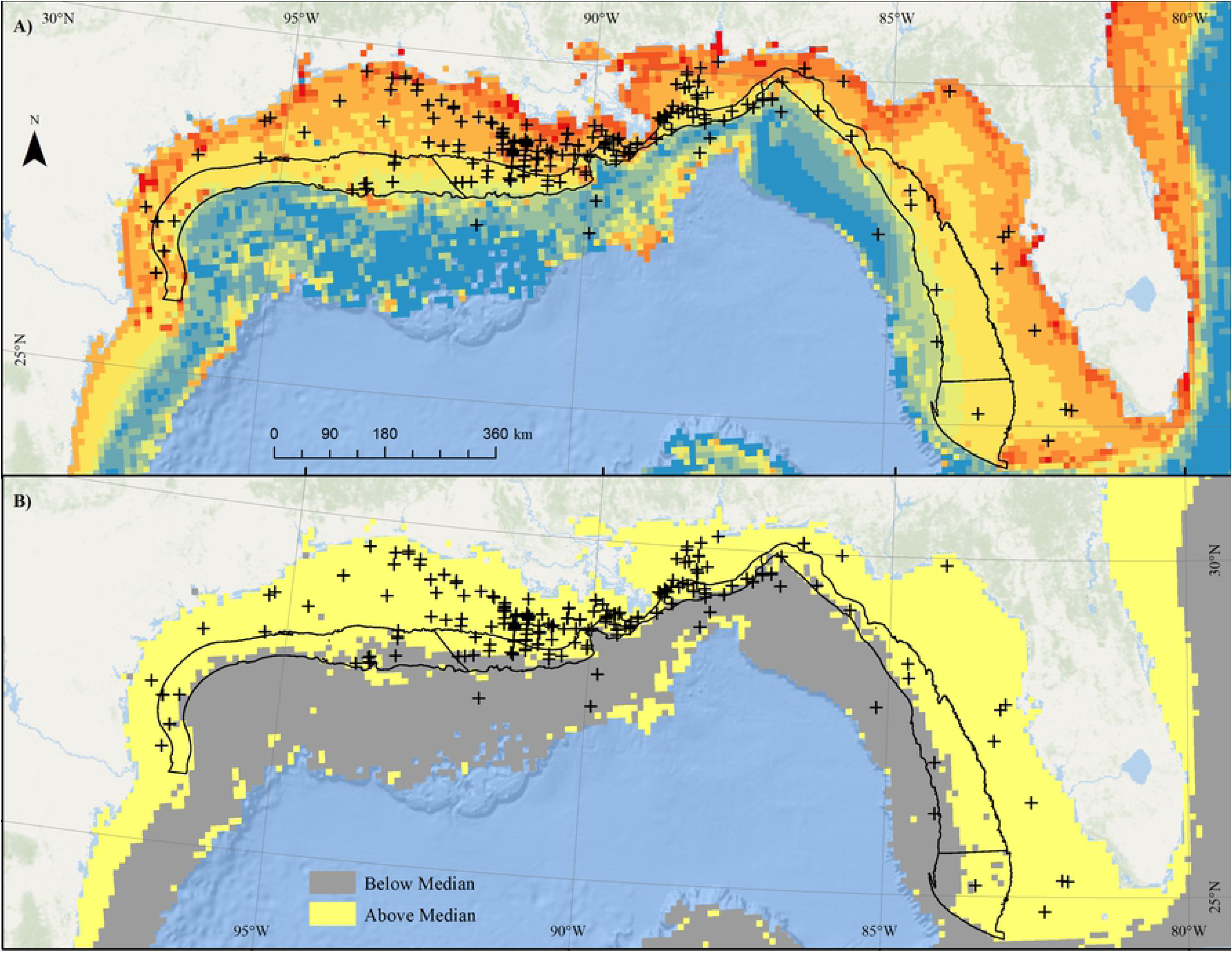
Map of A) maximum monthly predicted probability of Giant Manta Ray observation fit to environmental data from January 2003 to December 2019, overlaid with Giant Manta Ray sightings from 1925–2020, and B) areas falling above (yellow) and below (gray) median predicted probability of occurrence values. Black border denotes proposed aquaculture study areas. Map developed in ESRI ArcGIS 10.8.1. Basemap from Esri Ocean Basemap and its partners. Projection: UTM NAD83 Zone 17N.

### Combined Gulf of Mexico Protected Species Data Layer

In the Gulf of Mexico Atlas study areas, the Natural Resources submodel (**Figure 1**) contains a National Ocean Service sanctuaries data layer, consisting of only FGBNMS, and the combined protected species data layer described below. The FGBNMS layer is scored as a 0.5 within FGBNMS boundaries (e.g., unknown suitability) and a 1 in all other areas (i.e., “suitable” for aquaculture). Other areas not suitable for aquaculture [e.g., hard bottom and coral habitat areas of particular concern (CHAPCs)] are eliminated through the Constraints submodel from the AOA analysis after the mathematics are run to combine layers.

We compared four approaches to combining protected species data layers across species: 1) Product, 2) Geometric mean, 3) Arithmetic mean, and 4) Lowest scoring species in a given cell, using a custom R script. We combined protected species layers within each AOA subregion (i.e., West, Central, East, Southeast) and then merged the four subregions for comparison of scoring approaches, noting that although the combined protected species data layer can be compared between areas, Atlas study areas will be modeled independently and may not be comparable across Cumulative Suitability Models given potential differences in data types between areas.

Because the results of each Cumulative Suitability Model were used to identify the most suitable (top-ranked) clusters of cells to inform the development of preliminary alternatives of the AOA process, the relative ordering of scoring across cells was more important to the final outcome of the model than the actual scores within cells. We evaluated the outcomes of ranking protected species scores on a cell-by-cell basis, with ties ranked by minimum value similar to sports rankings (i.e., 1, 2, 3, 3, 4, 5, 6, 6, 7, etc.).

## Results

### Combining Protected Species Data Layers

Using a hypothetical distribution of three species with different distributions, status, and trend (**Figure 5**), we identified issues of concern with regards to applying the geometric and arithmetic mean approaches to ranked data layers (e.g., **Table 1**). For categorical scores (0/0.5/1) we anticipate the geometric mean works correctly because a 0 in a cell results in an overall score of 0, and more overlapping cells at 0.5 results in a cumulative lower score without a relative value system imposed. However, when multiple protected species layers overlap and a relative value scale is imposed on those layers, both the geometric and arithmetic mean methods result in final scores within individual cells that can be higher than the score for the species of greatest concern within the cell. For example, using the hypothetical distributions in **Figure 5**, the score for cells containing Species 1 is higher than 0.1 in both the arithmetic and geometric mean approaches, because those cells also contain Species 2 and/or Species 3. Also of concern, as more layers are incorporated with relatively limited spatial distributions for the species in question and the remaining areas are scored as 1s for that species, the geometric mean tends towards 1:

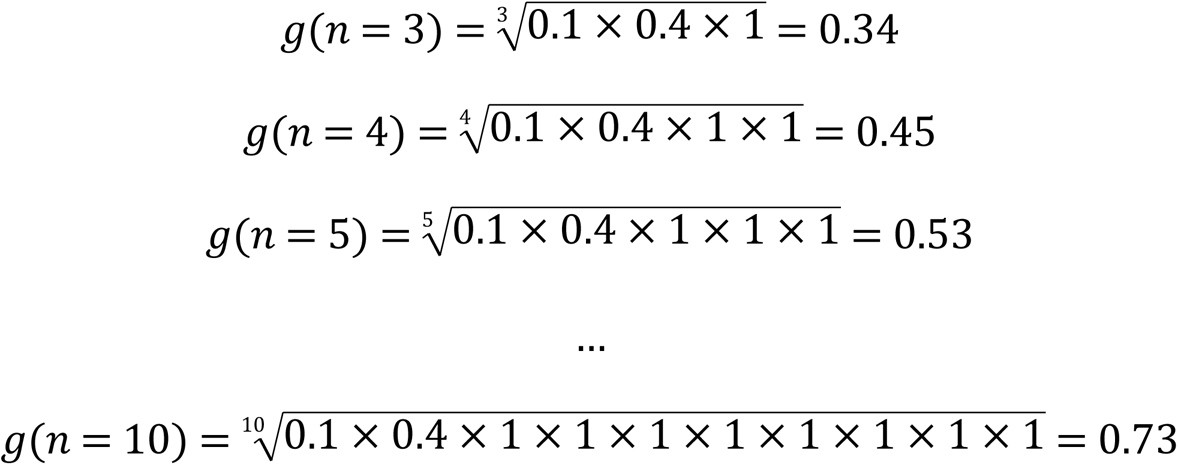

**Figure 5.**
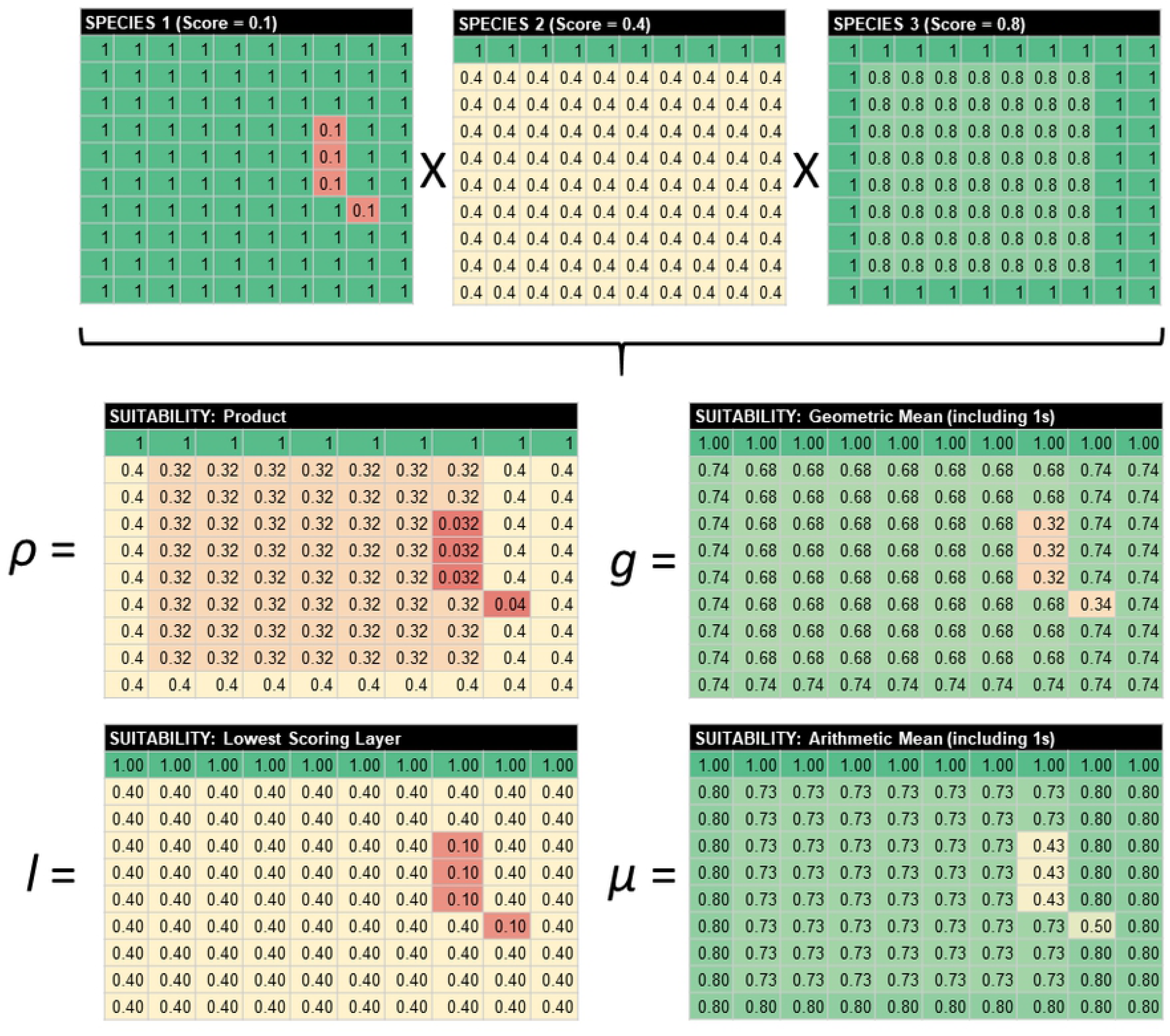
Hypothetical combination of three protected species data layers scored following Table 1, using four different approaches: Product (*ρ*), Geometric mean (g), Lowest Score (l), and Arithmetic mean (*μ*). Warmer colors denote areas of higher concern for protected species.

Therefore, both averaging approaches reduce the spread in the data with regards to differences between cells, and the geometric mean further compresses the data towards a score of 1 (i.e., no protected species in cell; area suitable for aquaculture). Only the Product approach generates scores below 0.1 (the minimum value for a single species in **Table 1**) when multiple species overlap. Mathematically, the averaging approaches can also result in different rank orders for cells as compared to the Product approach, depending on the scores for overlapping species layers (**Figure 5**).

### Case Study: Gulf of Mexico Aquaculture

Application of the four scoring approaches to protected species data layers in the Gulf of Mexico proposed aquaculture study areas provided additional support to the appropriateness of scoring using the Product approach. The resulting scores clearly show the greatest contrast between cells for the Product [median (M)=0.08, range (R)=0.000096–1) and Lowest (M=0.1, R=0.1–1) scoring layer approaches, as expressed by both visual spatial contrast between locations (**Figure S1**) and quantitative analysis of spread in the overall scoring (**Figure 6**). The arithmetic (M=0.81, R=0.39–1) and geometric mean (M= 0.73, R= 0.31–1) approaches generated the highest overall scores and least contrast between score (**Figure 6**).

**Figure 6.**
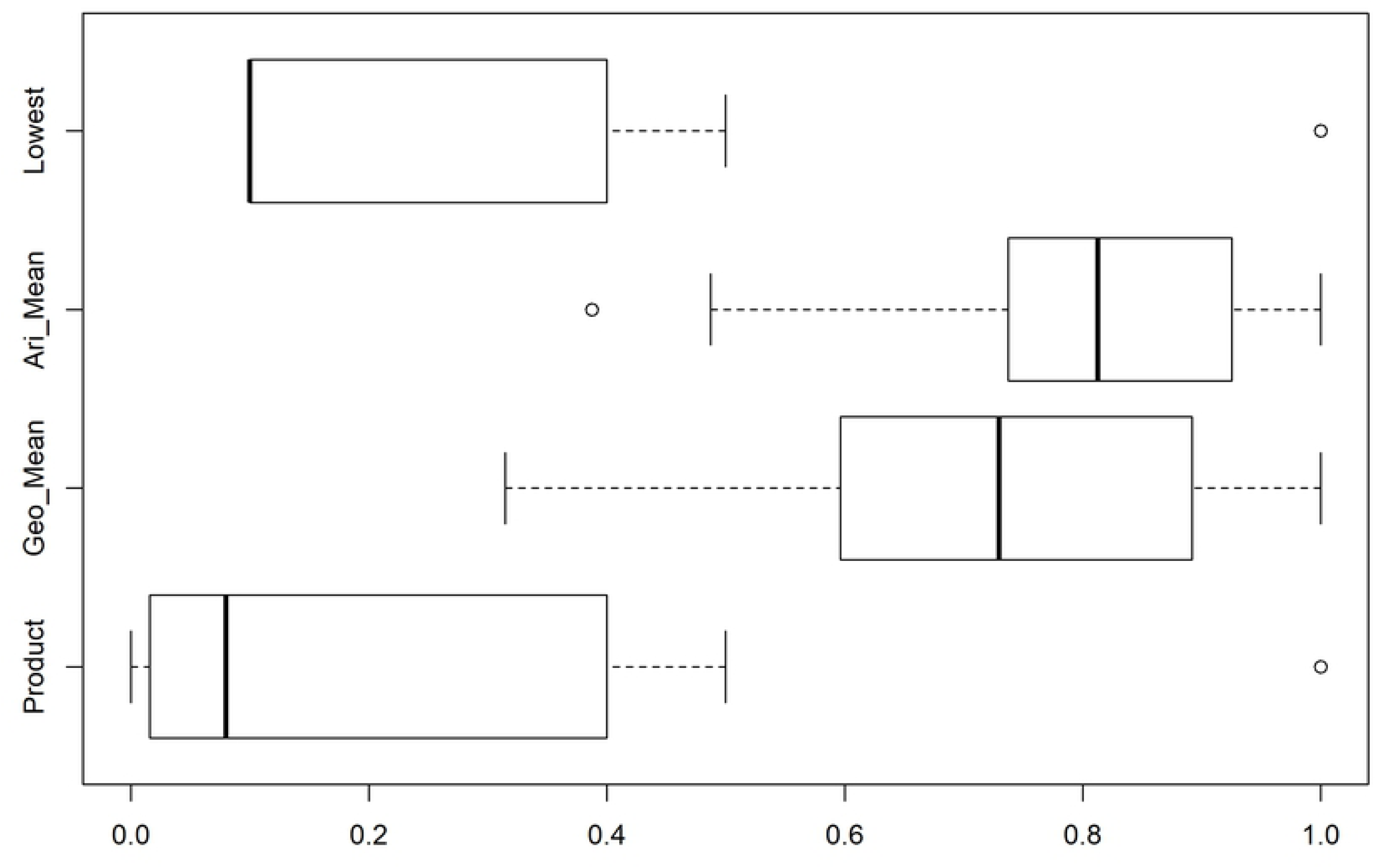
Comparison of vulnerability scores for protected species across the entire Gulf of Mexico study area under Product, Geometric mean, Arithmetic mean, and Lowest scoring layer methods. Lower scores denote greater vulnerability.

Rankings were inconsistent in 93% of cells when comparing between all four methods; the Lowest approach was the least consistent with regards to rankings of cells, presumably because it did not account for overlap of species in any way. Although the geographic distribution of ranks was relatively similar between the remaining three methods, the specific ranks for the averaging approaches were different from the Product approach in 62% of cells. As previously mentioned, the spread of the actual scores was substantially narrower in the averaging approaches, which may influence ultimate clustering outcomes, especially when combined with other data layers in the “Cumulative Suitability Model.” Unlike the majority of layers in the Gulf of Mexico Atlas, which are scored as 0/0.5/1 if they are categorical data, the protected species data layers have a relative value scale imposed (0.1–0.8), which, based on our evaluation, makes the Product approach more appropriate.

The final combined protected species data layers using the Product method indicated that high overlap of vulnerable protected species was a concern throughout the Central and East study areas, and also in the furthest offshore areas of the West proposed aquaculture study area (**Figure 7A**). The distribution of Rice’s Whale was the primary driver for these outcomes but was further informed by high use areas for several sea turtles and Giant Manta Rays. Similarly, the distribution of Smalltooth Sawfish was a primary driver for areas of high concern in the nearshore end of the Southeast study area (**Figure 7A**).

**Figure 7.**
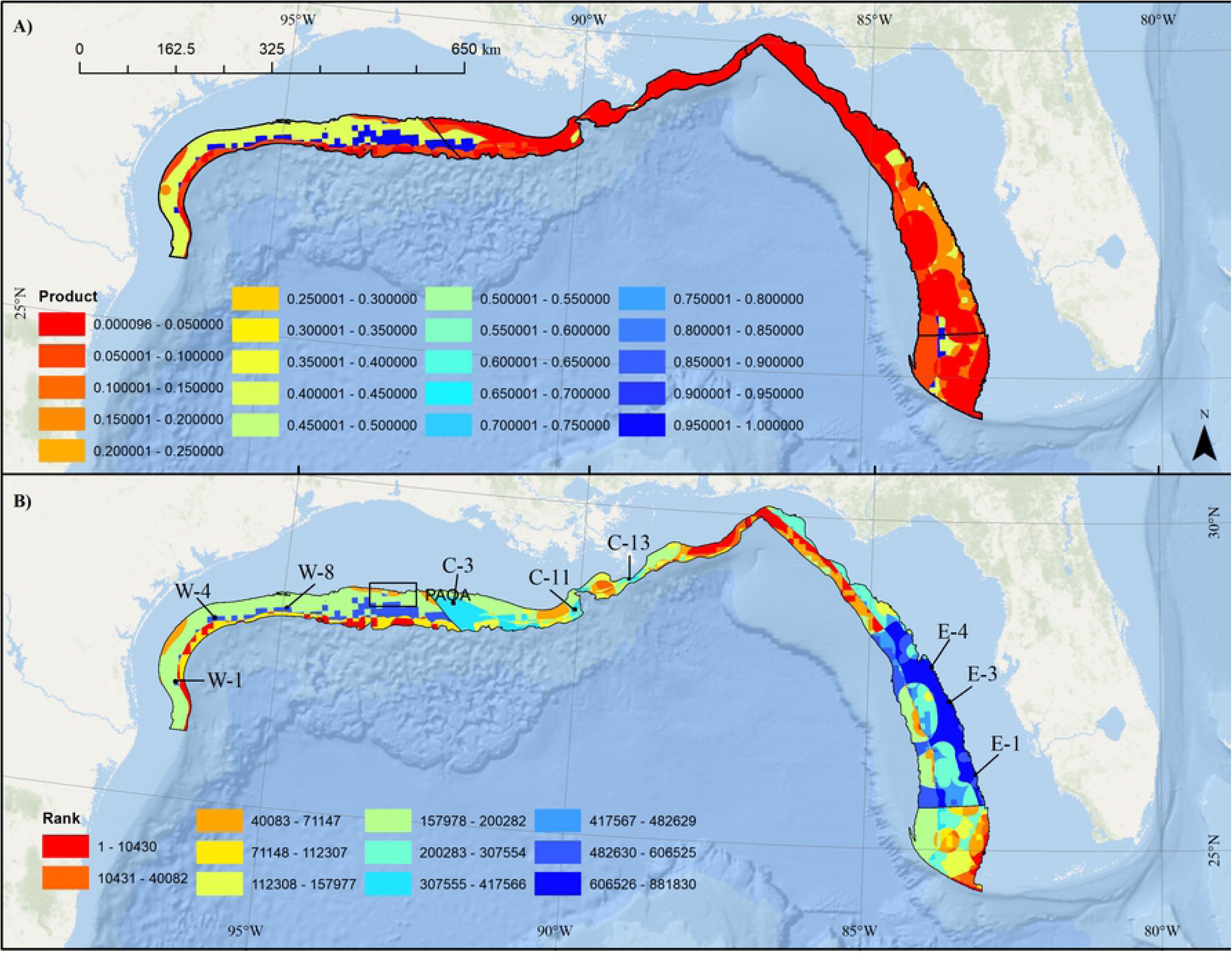
Combined Gulf of Mexico protected species data layer showing A) Product score across proposed aquaculture study areas (black) and B) Ranked scores using Product method relative to the nine final AOA options (i.e., W-1, W-4, W-8, etc.) within each of the four aquaculture study sub-areas (black). Layer generated by combining Rice’s Whale, five species of sea turtles, Smalltooth Sawfish, and Giant Manta Ray layers using the Product method. Warmer colors denote areas of relatively higher concern with regards to species status, population size, and trajectory. Map developed in ESRI ArcGIS 10.8.1. Basemap from Esri Ocean Basemap and its partners. Projection: UTM NAD83 Zone 17N.

The division of the study area into four separately considered subregions provided more contrast between cells based on within-subregion rankings (**Figure 7B**). In the West subregion, the area of least concern was in the middle of the proposed aquaculture study area off the border of Texas and Louisiana (**Figure 7B**). In the Central subregion, areas off western Louisiana were of lower concern than areas off Alabama and Florida (**Figure 7B**). In the Eastern subregion, only a few smaller nearshore areas were of lower concern, but still generally high concern relative to other areas in the Gulf of Mexico (**Figure 7**). In the Southeast subregion, shallower areas were of higher concern (**Figure 7B**).

Overall, the AOA suitability modeling results largely avoided areas of high importance for protected species (Riley et al. 2022). **Table 3** provides a listing of all protected species data layers included in the combined model. Of the 108 possible interactions between protected species and the top ten AOA alternatives, only 10 (9.2%) interactions occurred, eight of which were related to the Giant Manta Ray whose distribution is extensive. When excluding the Giant Manta Ray, only three (2.7%) interactions occurred (Loggerhead and Green Sea Turtle).

**Table. 3.**
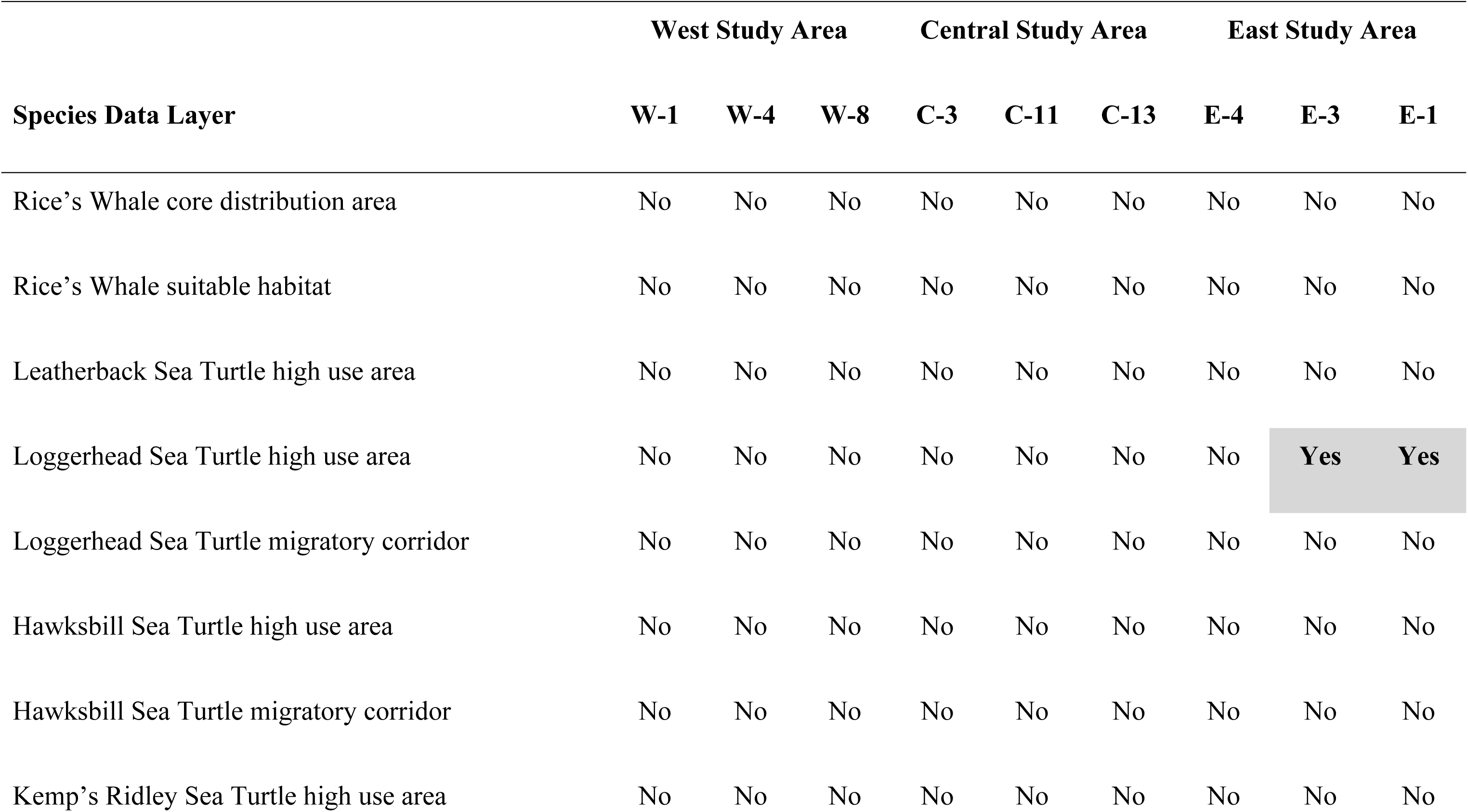
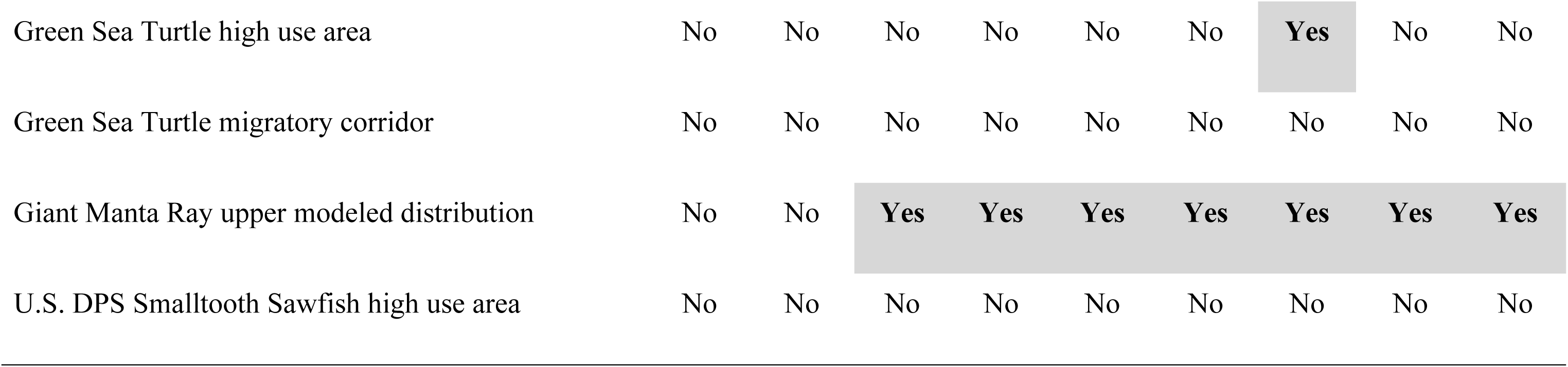
Top AOA options and interaction (yes/no) with protected species. Note the low level of interaction with protected species. Interactions in bold and shaded in gray.

|  | West Study Area |  |  | Central Study Area |  |  | East Study Area |  |  |
| --- | --- | --- | --- | --- | --- | --- | --- | --- | --- |
| Species Data Layer | W-1 | W-4 | W-8 | C-3 | C-11 | C-13 | E-4 | E-3 | E-1 |
| Rice's Whale core distribution area | No | No | No | No | No | No | No | No | No |
| Rice's Whale suitable habitat | No | No | No | No | No | No | No | No | No |
| Leatherback Sea Turtle high use area | No | No | No | No | No | No | No | No | No |
| Loggerhead Sea Turtle high use area | No | No | No | No | No | No | No | **Yes** | **Yes** |
| Loggerhead Sea Turtle migratory corridor | No | No | No | No | No | No | No | No | No |
| Hawksbill Sea Turtle high use area | No | No | No | No | No | No | No | No | No |
| Hawksbill Sea Turtle migratory corridor | No | No | No | No | No | No | No | No | No |
| Kemp's Ridley Sea Turtle high use area | No | No | No | No | No | No | No | No | No |
| Green Sea Turtle high use area | No | No | No | No | No | No | **Yes** | No | No |
| Green Sea Turtle migratory corridor | No | No | No | No | No | No | No | No | No |
| Giant Manta Ray upper modeled distribution | No | No | **Yes** | **Yes** | **Yes** | **Yes** | **Yes** | **Yes** | **Yes** |
| U.S. DPS Smalltooth Sawfish high use area | No | No | No | No | No | No | No | No | No |

## Discussion

Considerations for stationary protected resources, such as seagrass, corals, or marine protected areas, are often included in MSP models for aquaculture (Perez et al. 2005; Longdill et al. 2008; Lester et al. 2018). However, mobile or transient protected resources, such as marine mammals, are generally excluded from MSP. For mobile species, there is often uncertainty as to the impacts of ocean industries during the early planning stages, coupled with uncertainty regarding species distributions, sparse location data, or highly variable location data. However, in some cases, sufficient data are available for summary and integration into an MSP modeling approach. Early integration into planning processes reduces the likelihood of future conflict. Transparency about potential conflicts in the early planning stages can also help avoid contentious and time-consuming permitting decisions during the project design and implementation phase. In this study, we demonstrate how different forms of data for mobile protected resources may be summarized to inform distributions. We integrate across protected species scoring systems and layers using a generalized MSP approach that is portable across species and planning considerations. We identify the product methods as the most appropriate approach for combining data layers that have an internally consistent ranking scheme. Finally, we demonstrate the successful application of this approach to MSP for Gulf of Mexico aquaculture.

Spatial data from megafauna is used regularly by managers and policy makers to inform decisions regarding regulations and marine protected area boundaries (Hays et al. 2019). Tracking data and utilization density of Olive Ridley Sea Turtles in Pongara National Park, Gabon, was used for marine park and zone boundary designation (Dawson et al. 2017). In Augé et al. (2018), 36 species distribution layers of seabirds and pinnipeds were combined into a single composite megafauna layer using a weighted arithmetic mean to inform marine spatial planning around the Falkland Islands. In the presented case study here, eight data layers representing seven species were combined into a single composite layer for use within the suitability model.

MSP for aquaculture is generally performed using variants of a Multi-Criteria Decision Analysis, such as a suitability model; however, the methodology for how calculations are performed will vary by planning objective. In the Gulf of Mexico AOA MSP, an equally weighted geometric mean suitability model was determined to be the most representative for identifying suitable areas for general use aquaculture (Aguilar-Manjarrez 1996; Silva et al. 2011). Therefore, having equal weights ensures no differing expert opinions on weighting and no preference is given to one particular type of aquaculture or industry. Additionally, use of the geometric mean ensures that all variables contribute equally to the suitability score. The arithmetic mean assumes high suitability scores can compensate for low suitability scores, while the Product approach assumes high suitability scores cannot offset low suitability scores (Muñoz-Mas et al. 2012).

Both a hypothetical example and a case study application to siting for Gulf of Mexico aquaculture indicated that the Product approach was the most appropriate method for spatially combining overlapping protected species vulnerability scores (**Table 2**). The Product approach appropriately accounts for overlap between layers (**Figure 7A**), provides the correct ordering of the layers, and provides contrast between cells that should prove informative to the Atlas siting process (**Figure 7B**). Given consistency in methods and scoring, this final layer can be used to classify relative vulnerability for protected species both within and across the four sub-regions (i.e., West, Central, East, and Southeast) in the Gulf of Mexico Atlas. The final Atlas considered each subregion independently because other data sources could not be similarly compared. This resulted in equal numbers of AOA options from each sub-region besides the Southeast subregion, which had insurmountable spatial planning constraints with National Security activities. The use of subregions provided geographic parity across potential aquaculture options but did not reflect the final protected resources finding that conflicts were generally much higher in the northern and eastern Gulf of Mexico as compared to the western Gulf of Mexico.

This generalized scoring approach does not consider gear-specific risk associated with defined aquaculture activities. This was deliberate, as the Atlas was intended to inform aquaculture planning prior to having defined aquaculture activities proposed. Under the ESA, federal managers are required to avoid, minimize, or mitigate any federally-permitted activity that might jeopardize the continued existence or recovery of ESA-listed species. Our approach provides a transparent public-facing tool to avoid planning activities in areas with high overlap of vulnerable protected species. As such, the generalized protected species layer is more analogous to a map of consultation risk, with the final product helping to avoid conflict by reducing the likelihood of substantial investment in aquaculture design and development in areas where protected species overlap is high. A general principle of MSP is to preemptively avoid user conflicts through early engagement. Our method provides a transparent process to illustrate where any action requiring federal permitting would face numerous regulatory challenges due to high co-occurrence with vulnerable species.

Protected species data are often limited and subject to substantial bias and uncertainty, owing in part to the rarity of the study organisms as well as limitations on resources to study them. Our acoustic and satellite telemetry data were biased in focal species, life stages, and regions. Tagging locations are biased by tag location, size classes capable of carrying tags, and tag retention/track duration. Movements and space use of juvenile animals may differ substantially from those of adults. Fishery observer records were biased to fisheries with observer coverage, and to areas and time periods where fishing occurs. Fisheries may additionally only select for particular life stages, leading to bias in the overall picture of how the species is distributed.

Standardized aerial surveys were subject to both perception and availability bias that may be especially problematic for small, cryptic, or diving animals such as sea turtles and manta rays. The approach taken with Giant Manta Rays accounted for these biases and underlying aerial survey effort. Similar efforts to develop predictive models using aerial, ship, and PAM survey data are underway for most protected species. Those results should be incorporated into updates of the protected species data layer to facilitate adaptive management.

The application of this spatial modeling approach for protected resources has broad application across all ocean human uses and also for conservation. The novelty of this approach is that it provides the capability to synthesize across species and habitats providing a holistic perspective at a regional ocean scale and identifies “hotspots” where conservation is most critical when considering multiple species. While this approach was developed to inform development of aquaculture areas, a similar approach could be used for other development efforts including energy extraction, renewable energy (e.g., wind and marine hydrokinetic), shipping and transportation (e.g., development of new or modified shipping fairways), mining, dredging, and other types of activities that may impact protected resources.

This model provides a new approach for assessing the relative sensitivity of ocean spaces to development activities such as aquaculture at a regional scale. The application of this model will be useful when planning for changes of existing ocean industries, new pioneering ocean industries, and for conservation. Given the spatially and temporally dynamic nature of ocean space, frequent model updates will likely be required to ensure that the most accurate stock assessment data are being considered. Further development of environmentally-driven models similar to the Giant Manta Ray distribution model (Farmer et al. 2022) will provide additional spatiotemporal resolution and predictive capability. These types of models provide four major benefits: 1) they control for bias in observation effort, 2) they can be fit across a broad suite of environmental conditions observed across many years to provide a conservative estimate of long-term impacts, 3) they are adaptive to dynamic ocean conditions, and 4) they allow identification of spatiotemporal environmental windows where likelihood of species presence is low (Farmer et al. 2016). Environmental windows can be used to minimize adverse effects during the most impactful activities, such as construction. Generalized guidance will require revision for project activities that involve permanent or semi-permanent habitat change, which include most types of fixed-location operations.

The rapid development of wind energy in the U.S. and globally is one specific application where this modeling approach could better inform placement of wind farms on both regional and local scales (precision siting). Presently, wind energy development is being planned or underway across five regions of the U.S. (Northeast, Mid-Atlantic, Gulf of Mexico, South Atlantic, Pacific Coast) with additional regions being considered. Looking ahead to 2050, wind energy development constitutes one of the largest and most rapid consumers of ocean space with some reports suggesting that over 350,000 km^2^ of ocean space will be developed for the production of food and energy by 2050 compared to the approximately 40,000 km^2^ in 2018 (a 9-fold increase) (DNV 2021). Given the increasing pressure on ocean ecosystems globally, approaches such as the one presented here will be imperative to ensure conservation of our planet’s most sensitive species and habitats.

## Acknowledgements

*Disclaimer: The scientific results and conclusions, as well as any views or opinions expressed herein, are those of the author(s) and do not necessarily reflect those of NOAA or the Department of Commerce*.

Funding support provided by the NOAA National Centers for Coastal Ocean Science, the Department of Energy ARPA-E MARINER program, and the NMFS Office of Aquaculture. Aerial survey data for the Gulf of Mexico was collected during a study funded in partnership with the U.S. Department of the Interior, Bureau of Ocean Energy Management through Interagency Agreement M17PG00013 with the U.S. Department of Commerce, National Oceanic and Atmospheric Administration (NOAA).

## Appendices and Supplements

**Figure S1.**
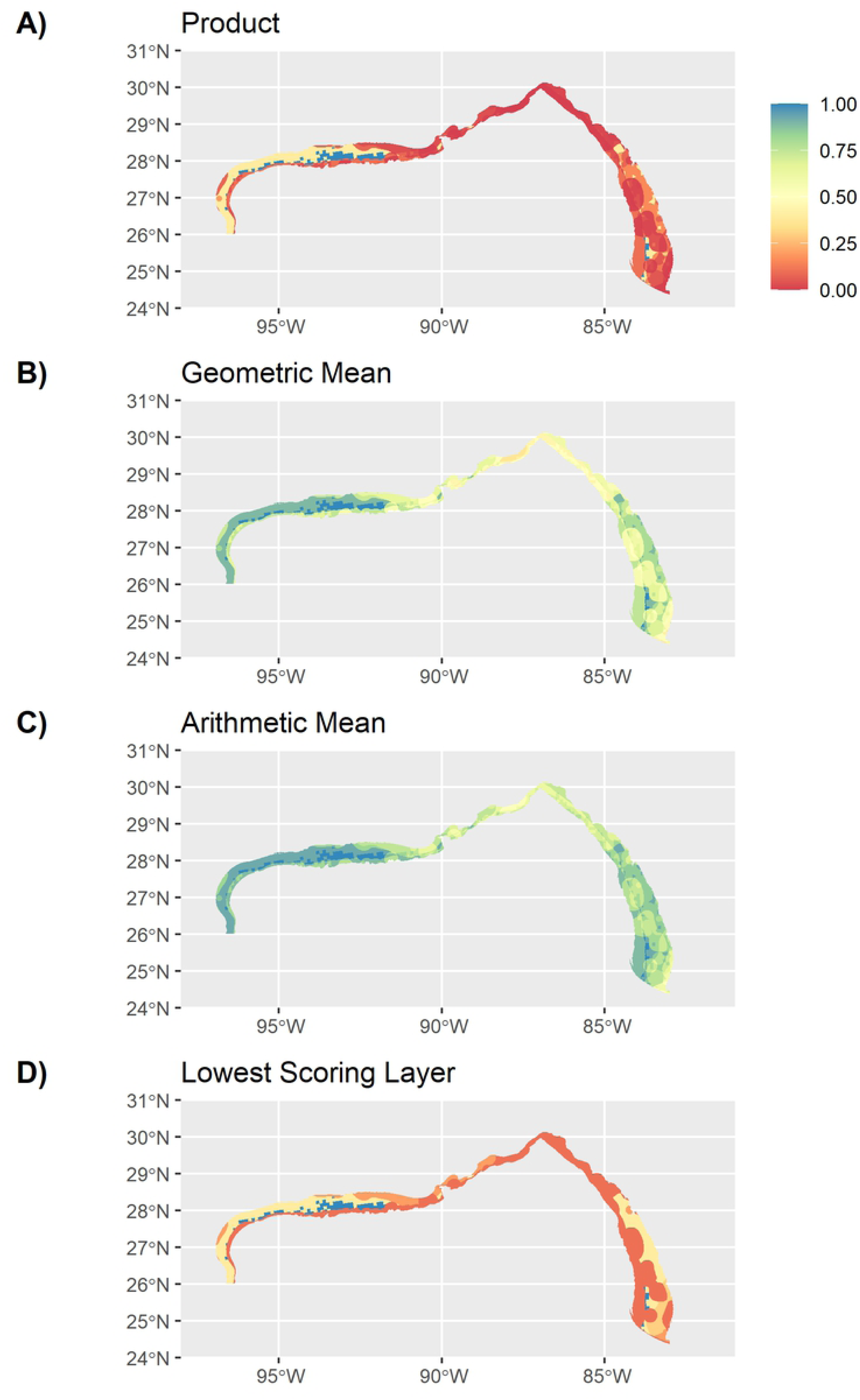
Comparison of overlapping protected resource data layer overall vulnerability scores combined using four different approaches. Warmer colors denote areas of greatest vulnerability.

